# Assessment of metagenomic sequencing and qPCR for detection of influenza D virus in bovine respiratory tract samples

**DOI:** 10.1101/2020.06.10.144782

**Authors:** Maodong Zhang, Yanyun Huang, Dale L. Godson, Champika Fernando, Trevor W. Alexander, Janet E. Hill

## Abstract

High throughput sequencing is currently revolutionizing the genomics field and providing new approaches to the detection and characterization of microorganisms. The objective of this study was to assess the detection of influenza D virus (IDV) in bovine respiratory tract samples using two sequencing platforms (MiSeq and Nanopore (GridION)), and species-specific qPCR. An IDV-specific qPCR was performed on 232 samples (116 nasal swabs and 116 tracheal washes) that had been previously subject to virome sequencing using MiSeq. Nanopore sequencing was performed on 19 samples positive for IDV by either MiSeq or qPCR. Nanopore sequence data was analyzed by two bioinformatics methods: What’s In My Pot (WIMP, on the EPI2ME platform), and an in-house developed analysis pipeline. The agreement of IDV detection between qPCR and MiSeq was 82.3%, between qPCR and Nanopore was 57.9% (in-house) and 84.2% (WIMP), and between MiSeq and Nanopore was 89.5% (in-house) and 73.7% (WIMP). IDV was detected by MiSeq in 14 of 17 IDV qPCR-positive samples with Cq (cycle quantification) values below 31, despite multiplexing 50 samples for sequencing. When qPCR was regarded as the gold standard, the sensitivity and specificity of MiSeq sequence detection were 28.3% and 98.9%, respectively. We conclude that both MiSeq and Nanopore sequencing are capable of detecting IDV in clinical specimens with a range of Cq values. Sensitivity may be further improved by optimizing sequence data analysis, improving virus enrichment, or reducing the degree of multiplexing.

## 1. Introduction

High throughput sequencing is currently revolutionizing the genomics field and providing new approaches to the detection and characterization of viruses. The utilization of metagenomic sequencing to elucidate genome sequences of viruses, particularly RNA viruses, directly from clinical samples offers several benefits. First, metagenomics enables identification and genomic characterization of unexpected viruses or even novel viruses either as primary pathogens or as co-infectants, without prior knowledge of their clinical significance [1]. Second, it eliminates the need for ongoing optimization of primers and/or probes for rapidly evolving or highly diverse RNA viruses [2]. Third, it facilitates routine surveillance and early detection of outbreaks of novel virus strains that are distinct from currently circulating strains. Finally, the development of portable sequencing devices creates the potential for timely identification of routine cases or outbreaks in the field [3,4]. Sequencing technology continues to evolve rapidly. With the capability of generating long reads, relatively lower set-up cost and portability, Oxford Nanopore sequencing has attracted increasing attention for its potential advantages in some circumstances over short-read sequencing technologies [5,6].

Metagenomic sequencing has been widely used for non-targeted detection of viruses and has been applied to identify several “new” viruses associated with bovine respiratory disease (BRD) [7-7]. In dairy cattle, bovine adenovirus 3 (BAdV3), bovine rhinitis A virus (BRAV) and influenza D virus (IDV) showed significant association with BRD [9]. In beef cattle, bovine rhinitis B virus (BRBV), BRAV and IDV showed statistical association with BRD [8,10]. Among all of the viruses detected by sequencing of the bovine respiratory tract metagenome, influenza D virus (IDV) has been identified as a common virus associated with BRD in both beef and dairy cattle, suggesting the potential contribution of IDV to BRD [8-8]. Influenza D virus (IDV) belongs to the Orthomyxoviridae, and is a single-stranded, enveloped, segmented and negative-sense RNA virus [11]. Since its discovery in swine in USA in 2011, IDV has been reported all over the world, and cattle are thought to be the natural host reservoir [10,12-12]. In addition to cattle and swine, IDV has been reported in sheep, goats, laboratory animals (ferrets and guinea pigs), and seropositivity has been detected in humans [14,16-16]. Concerns about interspecies transmission and potential zoonosis have been raised due to the high seroprevalence of IDV antibodies in people exposed to cattle [16].

Given the diversity of viruses now known to be associated with BRD and the potential for discovery of novel viruses like IDV, there is increasing interest in application of metagenomic sequencing for diagnostics, since screening for many individual viruses using targeted PCR assays quickly becomes logistically complex, expensive and time-consuming. The relative performance of metagenomic sequencing compared to PCR in terms of analytical sensitivity, however, has not been widely explored. In this study, IDV was used as a representative BRD-associated virus to examine the feasibility of using metagenomic sequencing for detection of viruses in clinical bovine respiratory samples. We compared results of long-read sequencing on the Oxford Nanopore GridION platform and previously generated Illumina MiSeq data [10] to the results of an IDV-specific qPCR. The objective was to assess IDV detection using all three approaches applied to a set of bovine respiratory tract samples containing a range of viral loads.

## 2. Materials and Methods

### 2.1. Ethics statement

The samples used in this study were collected as part of a previous study [10]. Collection of the samples was approved by the University of Calgary Veterinary Sciences Animal Care Committee (AC15 - 0109).

### 2.2. Sample preparation

The overall sample preparation workflow is shown in Figure 1a. Sample collection and preparation were described previously [10,19]. Briefly, paired nasal swabs (n = 116) and tracheal washes (n = 116) were collected from cattle with BRD and healthy controls from four different feedlots in Alberta, Canada between November 2015 and January 2016 [19]. The samples were centrifuged and supernatants were treated with DNase (Life Technologies, Carlsbad, CA) and RNase (Promega, Madison, WI), followed by extraction of viral nucleic acids using a commercial kit (QIAamp MinElute virus spin kit, Qiagen, Venlo, Netherlands). A portion (2.5 µl) of extracted total nucleic acids was used directly as a template for IDV qPCR and another portion (7 µl) used to generate cDNA for sequencing. The first strand was reverse transcribed with primer FR26RV-N using Superscript III enzyme (Life Technologies, Carlsbad, CA), followed by complementary strand synthesis using Sequenase polymerase (Affymetrix, Santa Clara, CA) as per manufacturer’s instructions [20]. Double-stranded DNA was purified using NucleoMag beads (Macherey-Nagel Inc., Bethlehem, PA) and subsequently subjected to random amplification with primer FR20RV prior to sequencing library preparation [20].

**Figure 1.**
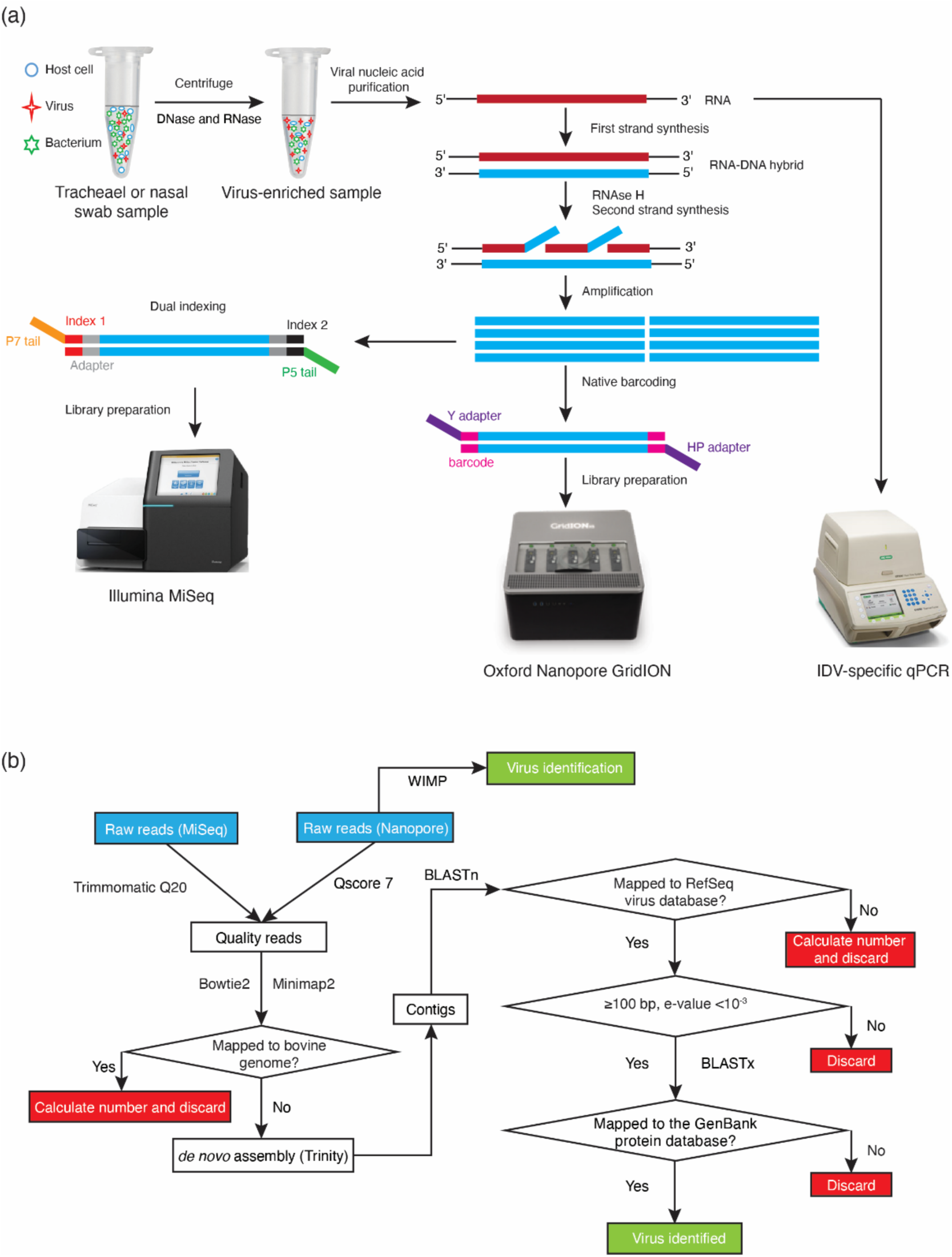
Workflow for GridION Nanopore sequencing, MiSeq sequencing, and qPCR of bovine respiratory tract samples. **(a)** The workflow illustrates both the sample and library preparation. Extracted RNA was used directly for qPCR, while DNA randomly amplified from the same extracts were used for MiSeq sequencing and GridION Nanopore sequencing; **(b)** Bioinformatic workflow to identify viruses in BRD samples. WIMP analysis was used only for Nanopore data. The remaining analysis was the same for data from both MiSeq and GridION Nanopore sequencing except Minimap2 was used instead of Bowtie2 in for host sequence subtraction.

### 2.3. qPCR confirmation and quantification

Quantitative real-time PCR for IDV was performed on the extracted total nucleic acids for each sample (total number of samples = 232) using previously described primers and probe specific for IDV [21]. The qPCR was carried out using AgPath-ID One-Step RT-PCR reagents in a total volume of 12.5 µl, which included 2.5µl template, 4 pM forward/reverse primers, 2 pM probe and 0.5 µl AmpliTaq Gold DNA polymerase in a Bio-Rad CFX 96 Real-Time Detection System (Bio-Rad, Hercules, CA). The following cycling conditions were used: reverse transcription phase at 48 °C for 30 min; initial activation phase at 95 °C for 10 min; 40 two-step cycles of denaturation at 95 °C for 15 s; and annealing and extension at 60 °C for 1 min. To obtain a positive control template, a DNA fragment corresponding to a 170 bp portion of the PB1 gene of IDV (accession number: JQ922306) was synthesized and inserted into pUC57-Amp vector (Bio Basic, Markham, ON). A 10-fold dilution series of the positive control plasmid was used to construct a standard curve to determine the efficiency of the PCR. All samples were tested in duplicate along with the standard curve and no template controls. Samples for which both of the duplicates gave a sigmoid amplification curve with a Cq (cycle quantification) value were considered positive.

### 2.4. GridION library preparation and sequencing

Nineteen samples that were IDV positive by either MiSeq virome sequencing or qPCR (with a range of representative Cq values) were conveniently selected for Nanopore sequencing. Three batches of six samples were multiplexed and run on individual flow cells, while one sample (sample 129) was run individually. The DNA used for GridION Nanopore library preparation was from the same randomly amplified DNA that was used for MiSeq sequencing (Figure. 1A). Ligation 1D sequencing kit SQK-LSK108 was used for library preparation. End-repair and dA-tailing were performed on randomly amplified DNA for each sample using NEBNext FFPE DNA repair mix and Ultra II End-prep enzyme mix (New England Biolabs, Ipswich, MA). After purification with AMPure XP beads (Beckman Coulter, Brea, CA), native barcode ligation using the EXP-NBD103 barcode kit (Oxford Nanopore Technologies, Oxford, UK) and Blunt/TA Ligase master mix (New England Biolabs, Ipswich, MA) was performed as per manufacturer’s instructions. The concentration of barcoded libraries was determined using a Qubit fluorometer (ThermoFisher Scientific, Waltham, MA) and subsequently equimolar amounts of each barcoded library (total amount = 1µg) were pooled, and adaptors were added using Quick T4 DNA Ligase (New England Biolabs, Ipswich, MA). Each pooled library (14.5 µl) was mixed with 35 µl Priming Buffer and 25.5 µl Loading Beads and loaded dropwise through the sample port into the flow cell (FLO-MIN106) as per manufacturer’s instructions. The MinKNOW platform QC check confirmed at least 800 available pores, and the High Accuracy Basecalling (HAC) Flip-flop model was applied.

### 2.5. Bioinformatic analysis

The workflow of bioinformatic analysis is illustrated in Figure 1b. Once Nanopore raw data were demultiplexed and trimmed using Porechop and passed the quality score (Qscore) 7, high quality reads were aligned to the bovine genome (BioProject Accessions PRJNA33843, PRJNA32899) using Minimap2, and unmapped reads (i.e. non-host derived reads) from each sample were de novo assembled using Trinity [22-22]. Assembled contigs were mapped to the virus Reference Sequence (RefSeq) database using BLASTn and virus-like contigs with a minimum alignment length of 100 bp and an expectation (e) value < 10^−3^ were further examined by BLASTx alignment to the GenBank non-redundant protein sequence database to confirm the nucleotide sequence-based identification and to remove any spurious matches [25]. The total number of viral reads was determined as previously described [10].

Quality filtered reads from the Nanopore sequencing were also uploaded to the EPI2ME platform for analysis with the WIMP (What’s in My Pot, version 2.3.7) application for taxonomic classification of reads.

## 3. Results

### 3.1. Comparison of IDV detection by MiSeq and qPCR

A total of 232 samples (116 nasal swabs and 116 tracheal washes) that had been sequenced using MiSeq previously (500 cycle V2 chemistry, libraries of 50 multiplexed samples) were tested by an IDV-specific qPCR (Figure 1a) [10]. The detection limit of the PCR was demonstrated to be 62.5 copies per reaction (data not shown). There were 53 IDV positive samples based on the qPCR and the range of Cq values was from 16.99 (6.25 × 10^7^ copies per reaction) to 39.46 (6.88 copies per reaction) with median Cq value being 34.07 (2.91 × 10^2^ copies per reaction). The agreement of IDV detection between qPCR and MiSeq was 82.8%. When qPCR was regarded as the gold standard, the sensitivity and specificity of MiSeq detection were 28.3% and 98.9%, respectively. IDV was detected by MiSeq in 14 of 17 IDV qPCR-positive samples with Cq values below 31 (Figure 2), when multiplexing 50 samples in the MiSeq flow cell. Only 1 of 36 IDV qPCR-positive samples with a Cq value above 31 was detected by MiSeq (Figure 2). Nineteen samples that were positive for IDV by qPCR or MiSeq, and that represented the full range of qPCR Cq values, were selected for further analysis by Nanopore sequencing.

**Figure 2.**
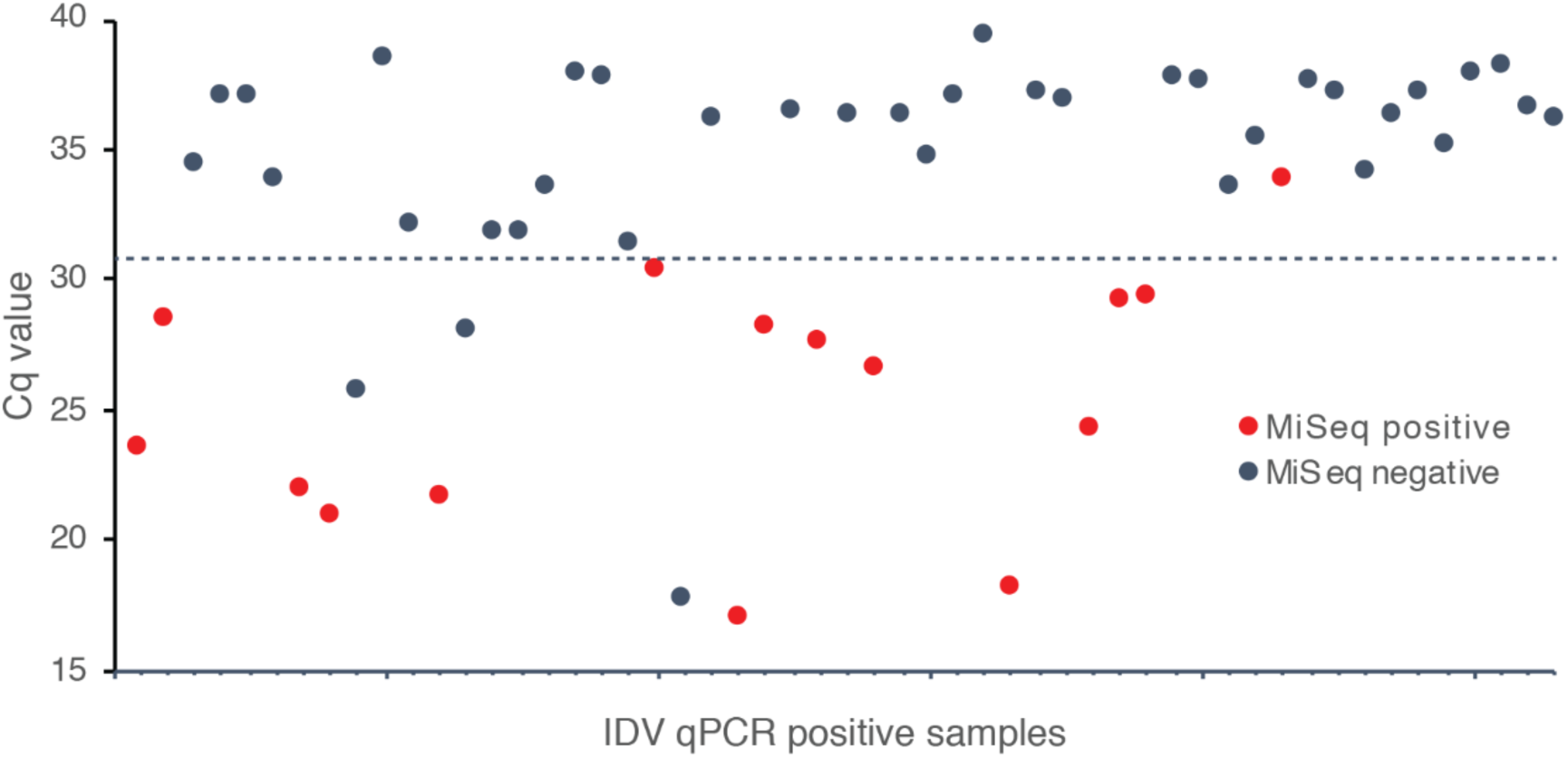
Cq values and MiSeq detection results for 53 samples positive by IDV qPCR. Samples positive for both qPCR and MiSeq are indicated by red dots. The majority of concurrent detection by MiSeq and qPCR occurred in samples with Cq values <31 (dotted line).

### 3.2. Comparison of Nanopore sequencing results to previously determined MiSeq data

A total of 82.7 million reads were obtained from MiSeq. When removed low-quality reads and host-derived reads, 33.6 million reads were remained. A total of 1.8 million high-quality viral reads were generated, accounting for 2.19% of the total reads obtained from MiSeq [10]. A total of 5.9 million Nanopore reads including unclassified (30.5%) and classified reads (69.5%) passed the quality filter (Qscore 7) using MinKNOW. After subtracting reads reported as “unclassified” by WIMP, a total of 0.41 million viral reads were obtained, accounting for 6.9% of total reads obtained (Figure 3). The proportion of viral reads per sample was 0.1% to 18.4% (Nanopore, WIMP analysis) compared to 0.03% to 3.1% for the previously generated MiSeq data; however, with both sequencing approaches, the majority of reads obtained were identified as host-derived or other (bacteria, fungi, unclassified) (Figure 3).

**Figure 3.**
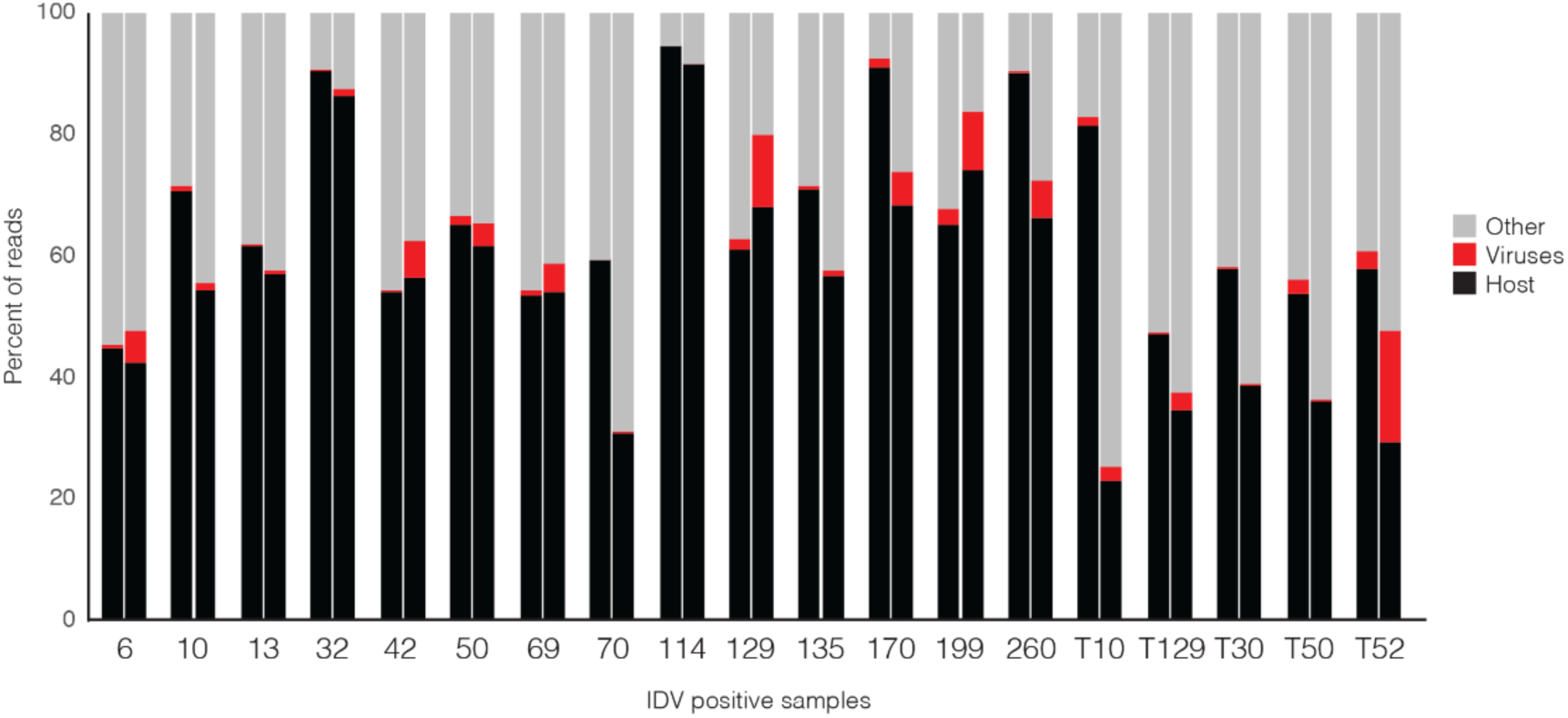
Proportions of reads corresponding to host, viruses or other taxa (bacteria, fungi, unclassified) from 19 IDV-positive samples sequenced using Nanopore sequencing (WIMP) (left bar in each pair) and MiSeq sequencing (right bar). Labels on the x-axis indicate individual specimens; tracheal samples are denoted by “T” before animal number.

In addition to WIMP classification of quality-filtered Nanopore reads, we also performed a *de novo* assembly of the Nanopore reads. The largest IDV contigs assembled for each sample from Nanopore data (using the in-house bioinformatics workflow, Figure 1b) were generally longer than those from MiSeq data and ranged from 626 to 2308 bp (Nanopore), and 249 to 1584 bp (MiSeq) (Table 1). The genome (or genome segment) coverage of each largest contig from each sample was from 10.53% to 95.31%. The proportion of Nanopore reads mapped to IDV for each sample by in-house analysis was higher than that from MiSeq except for sample T10 and T30. The proportion of IDV reads identified in the WIMP analysis of the Nanopore data, however, was generally comparable to that from the Nanopore (in-house) workflow (Table 1). The proportion of reads identified as IDV in Nanopore (WIMP), Nanopore (in-house) and MiSeq sequencing was generally extremely low (average 2.51%, 17.03% and 0.46%, respectively). As expected, approximately six times more reads were obtained for the individually sequenced sample 129 than for those from the multiplexed samples (Table 1). Sample 129 also had the lowest Cq value in the IDV qPCR (16.99, corresponding to 6.25 x 10^7^ copies per reaction) and the highest proportion of IDV reads in the metagenomic sequencing results (Nanopore-WIMP 14.69%, Nanopore-in-house 27.72%, MiSeq 1.48%) (Table 1).

**Table 1.** Summary of data from Nanopore, MiSeq and qPCR on detection of IDV for each individual sample.

| Sample* | Cq value | Copy number<br>(per reaction) | Number (%) of IDV reads |  |  | Largest IDV contig (bp) |  | Total input reads |  |
| --- | --- | --- | --- | --- | --- | --- | --- | --- | --- |
|  |  |  | Nanopore<br>(WIMP) | Nanopore<br>(In-house) | MiSeq | Nanopore<br>(In-house) | MiSeq | Nanopore | MiSeq |
| 129 | 16.99 | $6.25 \times 10^7$ | 321,638 (14.69) | 606,932 (27.72) | 2,182 (1.48) | 2030 | 951 | 2,188,805 | 147,341 |
| 114 | 17.53 | $1.31 \times 10^7$ | 8 (<0.01) | ND | ND | N/A | N/A | 335,559 | 136,961 |
| 50 | 20.71 | $1.25 \times 10^6$ | 1,088 (5.74) | 1625 (8.57) | 162 (1.1) | 1161 | 498 | 18,966 | 14,719 |
| 69 | 21.69 | $4.48 \times 10^5$ | 944 (7.83) | 2287 (18.97) | 1,812 (0.69) | 980 | 842 | 12,053 | 263,262 |
| 42 | 22.04 | $2.86 \times 10^5$ | 608 (0.34) | 584 (0.33) | 26 (0.07) | 986 | 499 | 179,559 | 37,560 |
| 6 | 23.33 | $2.16 \times 10^5$ | 281 (1.44) | 350 (1.8) | 183 (0.08) | 1034 | 522 | 19,512 | 228,875 |
| 10 | 24.09 | $1.31 \times 10^5$ | 1,656 (1.17) | 1650 (1.11) | 87 (0.2) | 1359 | 485 | 148,381 | 44,433 |
| 199 | 26.01 | $4.64 \times 10^4$ | 4,211 (10.99) | 10861 (28.34) | 3,250 (1.49) | 2308 | 552 | 38,329 | 218,800 |
| T50 | 26.76 | $2.93 \times 10^4$ | 3 (<0.01) | ND | ND | N/A | N/A | 43,193 | 1,256,918 |
| T129 | 28.20 | $8.00 \times 10^3$ | 403 (0.23) | 502 (0.28) | 34 (<0.01) | 843 | 327 | 177,706 | 444,979 |
| 70 | 28.71 | $5.70 \times 10^3$ | 2 (0.02) | ND | ND | N/A | N/A | 12,211 | 160,704 |
| T10 | 29.15 | $5.50 \times 10^3$ | 29 (0.54) | ND | 455 (0.03) | N/A | 913 | 5,413 | 1,415,256 |
| 13 | 29.22 | $4.15 \times 10^3$ | 116 (0.10) | 152 (0.13) | 48 (0.04) | 626 | 470 | 119,073 | 132,847 |
| 32 | 33.60 | $2.18 \times 10^2$ | ND | ND | ND | N/A | N/A | 275,387 | 32,312 |
| 170 | 35.62 | $8.25 \times 10^1$ | 1,080 (2.03) | 1991 (3.75) | 8,167 (0.74) | 903 | 1,584 | 53,131 | 1,100,167 |
| 135 | 36.51 | $4.88 \times 10^1$ | 10 (<0.01) | ND | ND | N/A | N/A | 142,737 | 154,220 |
| 260 | 39.46 | 6.88 | 9 (<0.01) | ND | ND | N/A | N/A | 164,607 | 959,935 |
| T30 | ND | - | 3 (<0.01) | ND | 2 (<0.01) | N/A | 249 | 139,037 | 1,156,213 |
| T52 | ND | - | 51 (0.1) | 52 (0.1) | 497 (0.05) | 1616 | 1,341 | 51,180 | 930,628 |

Cq = quantification cycle; ND = not detected; WIMP = What's In My Pot; bp = basepair
\*Samples beginning with T are tracheal, all others are nasal swabs.

### 3.3. Comparison of IDV detection by qPCR, MiSeq and Nanopore sequencing

The 19 samples selected for sequencing on the Nanopore GridION platform represented a range of IDV concentrations based on qPCR of 6.88 to 6.25×10^7^ genome copies per reaction, corresponding to Cq values ranging from 39.46 to as low as 16.99 (Table 1). The agreement of IDV detection between qPCR and Nanopore was 57.9% (in-house) and 84.2% (WIMP), and that between MiSeq and Nanopore was 89.5% (in-house) and 73.7% (WIMP). IDV was detected in the Nanopore data from all but one (18/19) of the IDV-positive samples when reads were classified using the WIMP application, but this proportion dropped to 11/19 when the in-house read assembly workflow was used. For most (7/8) of the samples with disparate results, 10 or fewer IDV reads were identified in the WIMP analysis. The exception was sample T10 with 29 IDV reads.

In order to explore qualitatively whether detection of other viruses in addition to IDV was comparable between the two metagenomic sequencing platforms, we compared the complete lists of viruses detected by MiSeq or Nanopore in the 19 IDV-positive samples. The number of different viruses detected in each sample varied from none to a maximum of four. The proportion of samples with perfect agreement between MiSeq and Nanopore (in-house) was 52.6%, by MiSeq and Nanopore (WIMP) was 47.4%; and by Nanopore (in-house) and Nanopore (WIMP) was 36.8% (Supplementary Table 1).

## 4. Discussion

Metagenomic sequencing is transforming routine detection of viruses from traditional cell culture, antibody-antigen techniques and qPCR to detection of viruses in a target-independent manner. Sequencing approaches have now been widely applied for detection of known and novel agents in various types of clinical specimens in both human and veterinary medicine [26-26]. The potential usefulness of viral metagenomics for virus surveillance and diagnostics is still in debate due to its performance relative to the gold-standard method of real-time qPCR routinely employed in diagnostic laboratories [3]. A recent assessment of the performance of Nanopore, MiSeq and qPCR for detection of chikungunya and dengue viruses in serum or plasma samples with relatively high viral loads (Cq values from 14 to 32) demonstrated 100% agreement among these methods [1]. In this investigation, however, a maximum of 16 samples were multiplexed and sequenced using MiSeq, and each sample was sequenced individually on Nanopore [1]. This low degree of multiplexing translates to high analytical sensitivity, but correspondingly makes these technologies relatively more expensive per sample than expected, and decreases the potential application for routine diagnostics. In our current study, we performed further exploration to assess the performance of metagenomic sequencing approaches with a higher degree of multiplexing of clinical samples in both MiSeq and Nanopore sequencing. IDV presented an excellent target for this comparison given its association with BRD in beef and dairy cattle, and the availability of specimens with a wider range of viral loads than has been included in previous investigations [19,20,32].

Different viral extraction kits have been demonstrated to have variable extraction efficiencies for different viruses in respiratory clinical samples [29]. The QIAamp MinElute Virus Spin Kit (MVSK) has been found to be generally applicable for isolating nucleic acid for qPCR or metagenomic virus identification of adenovirus, influenza virus A, human parainfluenza virus 3, human coronavirus OC43 and human metapneumovirus in respiratory clinical samples [29]. In our current study, the original nucleic acids extracted with the MVSK were used for both qPCR and metagenomic sequencing, eliminating the influence of different extraction methods and kits on our results (Figure 1a).

The IDV-specific qPCR assay detected its target in 22.8% (53/232) of specimens. While the majority of IDV positive samples with Cq value below 31 were detected by MiSeq, only 1/36 samples with a Cq above this threshold were positive by sequencing (Figure 2). These results demonstrate that for samples where the viral load exceeds 6.25 × 10^2^ even a relatively modest MiSeq sequencing effort (50 samples multiplexed in a single flow cell) is sufficient to detect the virus. The agreement between qPCR and Nanopore of 57.9% (in-house) and 84.2% (WIMP) demonstrated that relatively modest Nanopore sequencing effort (6 samples multiplexed) is also sufficient to detect the virus. The results from Nanopore sequencing (in-house), however, showed no consistent relationship between viral load and detection by sequencing; furthermore, no consistent relationship between viral load and proportion of viral reads was observed in either MiSeq or Nanopore sequencing (Table 1). For example, the two IDV positive samples 199 and 10 had Cq values of 26.01 and 24.09, respectively; however, sample 199 had a higher proportion of IDV sequence reads in Nanopore and MiSeq than sample 10 (Table 1).

There are several possible explanations for differences in both the proportion of IDV reads and total viral reads detected in each sample by MiSeq and Nanopore. First, variation in the amounts of DNA used for sequencing library preparation for the two sequencing platforms may play an important role. Second, the abundance of virus relative to host or bacterial genetic material is a critical determinant of the detection threshold of metagenomic sequencing. A greater proportional abundance of a virus increases the chance that it will be detected by sequencing and improves the genome coverage obtained. Therefore, virus enrichment is commonly applied to clinical samples and enrichment methods such as those used in this study (a combination of centrifugation and nuclease-treatment) should lead to removal of bacteria and host cells, thus improving virus detection [30]. Virus propagation in cell culture is a less appealing method for virus enrichment since it is time-consuming, requires specific expertise and creates the potential for introduction of mutations [31]. Reduction of the degree of multiplexing of samples is an alternative way to improve virus detection, but there is a corresponding increase in cost per sample and a corresponding reduction in throughput that are undesirable in research or clinical diagnostic settings. Reduction of the degree of multiplexing of samples also reduces the chances of cross-barcode contamination because barcode reagents are susceptible to cross-contamination [32].

Bioinformatic analysis in metagenomic sequencing remains challenging but is crucial for accurate identification of diagnostic targets. We used the comparable pipeline to analyze both data from MiSeq and Nanopore sequencing (in-house), which showed the exciting feasibility of metagenomic viral whole-genome-sequencing using both Nanopore and MiSeq technology with the assembled contigs covering from 10 to 95% of each IDV genome segment. Although de novo assembly was performed on Nanopore sequencing data, for the majority of the samples, the length of the largest contig was that of one single read. Skipping the assembly step in bioinformatic analysis of Nanopore data could provide an advantage for timely identification of potential pathogens. The long reads of Nanopore sequencing is thought to provide good confidence for species level identification, but the low coverage combined with the error rates of this platform preclude its use for strain-level resolution [33].

The detection rates of IDV, the number or proportion of IDV reads (Table 1) and other viruses detected (Supplementary Table 1) from Nanopore (WIMP) and Nanopore (in-house) were different, which demonstrates that bioinformatic analysis affects the results of virus detection. Taxonomic classification in WIMP is based on Centrifuge [34], which compares query sequences to a (undescribed) reference database with a high speed and space-optimized k-mer-based algorithm. For Nanopore (in-house) and MiSeq analysis, BLAST [35] was used to compare assembled contig sequences to the NCBI RefSeq virus database, which is a more computationally intense process that produces more detailed results. The identification of a match using Centrifuge is based on probabilities of particular k-mer combinations occurring in the query and reference and not a consideration of the entire query sequence, thus increasing the possibility of false positives [34]. In contrast, Trinity assembly and then BLAST search against a reference database could lead to false negatives if the particular target sequence is very rare [36]. If there are very few reads derived from some component of the metagenome, these reads may not be included in the assembly since there is insufficient “evidence” to support building contigs from them [35,36]. The current lack of definition of the reference database or the ability to use custom databases with WIMP make this approach inappropriate for clinical diagnostic applications due to the difficulty of validating such approaches. Our results provide an illustration of the profound effects that post-sequencing analysis can have on results, and the trade-offs associated with each choice. Selection of the most appropriate analysis pipeline must consider the sequencing platform, as well as tolerance for false negatives and false positives, logistical considerations, and the required taxonomic resolution.

Analytical sensitivity is currently one of the main limitations of metagenomics. In this study, IDV was detected by MiSeq sequencing in specimens with qPCR Cq value as high as 35.62 when 50 samples were multiplexed in comparison to a maximum Cq value of 39.46 using Nanopore with multiplexing of 6 samples. For the IDV positive samples with low virus loads (e.g. sample 32), targeted qPCR may be preferable given its higher analytical sensitivity. Interestingly, we observed two samples that were IDV positive by both Nanopore (WIMP) and MiSeq but negative by qPCR (Samples T30 and T52, Table 1). These cases illustrate a potential advantage of metagenomic sequencing compared to qPCR since a likely explanation for this observation is that these specimens contained strain variants of IDV that were not detected by the qPCR assay. We were unable to determine if this was the case since the IDV sequence reads did not cover the region of the genome targeted by the species-specific qPCR assay. Targeted PCR assays for rapidly evolving RNA viruses require ongoing performance monitoring, and optimization of primers and probes [2]. No single method is suitable for application for all pathogens or specimen types, and each one has advantages in different circumstances.

Taken together our results demonstrate the potential of metagenomic sequencing on the Illumina MiSeq and Oxford Nanopore platforms for detection of viruses, including IDV, in clinical samples from naturally infected animals with a wide range of viral loads. While application of these approaches to screening animal populations or infectious disease research is feasible, their deployment for routine virology diagnostics in clinical settings will require additional research, laboratory and bioinformatic method development, and performance evaluation. Selection of appropriate methods will continue to require careful consideration of the numerous trade-offs that confront practitioners at each step of the investigation.

## Acknowledgments

We thank Anju Tumber (Prairie Diagnostic Services, Inc.) for reagent purchasing and other logistics. We are grateful for Kara Toews, Anatoliy Trokhymchuk, Kazal Krishna Ghosh (PDS) for technical support.

## Author Contributions

Conceptualization, M.D.Z, J.E.H and Y.Y.H.; methodology, M.D.Z.; software, M.D.Z and J.E.H.; validation, M.D.Z., Y.Y.H. and J.E.H.; formal analysis, M.D.Z.; investigation, Y.Y.H. and T.W.A.; resources, T.W.A.; data curation, M.D.Z., D.L.G. and J.E.H.; writing—original draft preparation, M.D.Z.; writing—review and editing, J.E.H., C.F., Y.Y.H. and D.L.G.; visualization, M.D.Z. and J.E.H.; supervision, Y.Y.H. and J.E.H.; project administration, Y.Y.H.; funding acquisition, Y.Y.H.. All authors have read and agreed to the published version of the manuscript.

---

**Supplementary Table 1.** Summary of viruses detected by Nanopore and MiSeq sequencing.

| **Sample\*** | **MiSeq** | **Nanopore (In-house)** | **Nanopore (WIMP)** |
| --- | --- | --- | --- |
| 6 | IDV | IDV | IDV |
| 10 | BRSV, BRBV, IDV | BRSV, BRBV, IDV | BRSV, BRBV, IDV |
| 32 | BNV | BNV | BNV |
| 50 | BRBV, IDV | BRBV, IDV | BRBV, IDV |
| 69 | IDV | IDV | IDV |
| 129 | BRBV, IDV | BRBV, IDV | BRBV, IDV |
| 170 | IDV | IDV | IDV |
| 199 | BRSV, IDV, UBPV6 | BRSV, IDV, BPIV3 | BRSV, IDV, UBPV6 |
| T10 | BRSV, IDV | BRSV | BRSV, IDV |
| 13 | IDV | IDV | BRSV, BRBV, IDV |
| T52 | BRBV, IDV | IDV | BNV, IDV |
| 42 | BCV, IDV | BCV, IDV, BRSV | BCV, IDV, BRSV, BNV |
| 114 | BRBV, UTPV1 | ND | BRSV, BRBV, IDV, UTPV1 |
| T30 | BRBV, IDV | BRBV | BRBV, IDV, BNV |
| T129 | BAV, IDV | BAV, IDV | BAV, IDV, BNV |
| 260 | EVE, UTPV1, UBPV6 | ND | BRSV, BRBV, IDV |
| 70 | BRAV | ND | BRSV, BRBV, IDV |
| 135 | BNV | ND | BRSV, BRBV, IDV |
| T50 | ND | ND | BNV, IDV |

**Footnote.** BCV: bovine coronavirus; IDV: influenza D virus; BRBV: bovine rhinitis B virus; BRAV: bovine rhinitis A virus; BRSV: bovine respiratory syncytial virus; BPIV3: bovine parainfluenza virus 3; EVE: enterovirus E; UTPV1: ungulate tetraparvovirus 1; UBPV6: ungulate bocaparvovirus 6; BNV: bovine nidovirus; BAV: bovine astrovirus; \*Samples beginning with T are tracheal, all others are nasal swabs, ND – not detected. WIMP – What’s in my pot.

